# Full-length native CGRP neuropeptide and its stable analogue SAX, but not CGRP peptide fragments, inhibit mucosal HIV-1 transmission

**DOI:** 10.1101/2021.03.08.434421

**Authors:** Morgane Bomsel, Anette Sams, Emmanuel Cohen, Alexis Sennepin, Gabriel Siracusano, Francesca Sanvito, Lars Edvinsson, Lucia Lopalco, Yonatan Ganor

## Abstract

The vasodilator neuropeptide calcitonin gene-related peptide (CGRP) plays both detrimental and protective roles in different pathologies. CGRP is also an essential component of the neuro-immune dialogue between nociceptors and mucosal immune cells. We previously reported that CGRP strongly inhibits human immunodeficiency virus type 1 (HIV-1) infection, by suprresing Langerhans cells (LCs)-mediated HIV-1 transfer to T-cells during trans-infection. To understand the requirements for CGRP receptor (CGRP-R) activation during inhibition of HIV-1 transmission, we here investigated the anti-HIV-1 activities of full-length native CGRP and its recently developed stable analogue SAX, as well as several CGRP peptide fragments containing its binding C-terminal and activating N-terminal regions. We show that SAX significantly inhibits LCs-mediated HIV-1 trans-infection but with lower potency than CGRP, while all CGRP peptide fragments tested have no effect. In addition, CGRP readily enters the epithelial compartment of mucosal tissues and does not modify the distribution and density of mucosal immune cells. *In-vivo*, a single CGRP treatment in humanized mice, before vaginal challenge with high-dose HIV-1, restricts the increase in plasma viral load and maintains higher CD4+ T-cell counts. Together, our results call for the optimization and design of CGRP analogues and agonists, which could be harnessed for prevention of mucosal HIV-1 transmission.

## Introduction

CGRP is a 37 amino acids potent vasodilator neuropeptide secreted from peripheral sensory nerves, which plays important physiological and pathophysiological roles (1). CGRP-R antagonism is clinically effective in migraine and several CGRP-R antagonistic molecules have already entered the market. Yet, CGRP-mediated vasodilation is protective, at least during hypertension and cardiovascular complications (1), and the first metabolically stable CGRP agonistic analogue SAX was only recently developed for preclinical studies. Although SAX has a longer halflife and lower potency than CGRP, both molecules share vascular pharmacological effects *in-vitro* (2) and *in-vivo* (3).

Native CGRP also directly modulates immune function in a vasodilator-independent manner, as part of the neuro-immune dialogue between CGRP-secreting mucosa-innervating nociceptors and resident immune cells (4). For instance, nociceptors associate with LCs and CGRP shifts LCs-mediated antigen presentation and cytokine secretion from Th1 to Th2/Th17 (5). We previously reported that LCs are the early cellular targets of HIV-1 upon its mucosal entry in the inner foreskin, which subsequently transfer infectious virus to CD4+ T-cells during trans-infection (6). We further discovered that CGRP activates the CGRP-R in LCs and modulates a multitude of cellular and molecular processes, resulting in significant inhibition of LCs-mediated HIV-1 trans-infection (7–9). Hence, CGRP analogues and agonists might be useful for prevention of mucosal HIV-1 transmission.

The N-terminus (residues 1-7, containing a disulphide bond between the cysteines at positions 2 and 7) and amidated C-terminus (residues 27-37) of CGRP interact independently with CGRP-R in a two-domain model, where the C-terminus first binds the receptor, facilitating subsequent binding and activation by the N-terminus (10). The N-terminal disulphide loop is crucial for agonistic activity, as the peptide fragment CGRP_8-37_ is an antagonist and as several N-terminal peptide fragments are agonistic and anti-hypertensive, although with decreased potency (11). Other CGRP peptide fragments, containing constrained N-terminus (i.e. truncated loop with only three residues) and/or introduced disulphide bridge in the C-terminus, yield analogues with affinities comparable to native CGRP (12).

Herein, to elucidate the requirements of CGRP-R activation for inhibition of HIV-1 transmission, we compared the anti-HIV-1 activities of full-length native CGRP, its analogue SAX, and several CGRP N-terminal fragments and N+C-terminal bivalent fragments. We also tested whether testing CGRP inhibits HIV-1 transmission *in-vivo*.

## Results and Discussion

### CGRP and SAX, but not CGRP peptide fragments, inhibit LCs-mediated HIV-1 trans-infection *in-vitro*

We used monocyte-derived LCs (MDLCs) to compare HIV-1 trans-infection inhibition by CGRP, SAX and CGRP peptide fragments (Figure 1A). Both CGRP and SAX significantly inhibited HIV-1 trans-infection in a dose-dependent manner, although SAX had lower potency with IC_50_ of 1.4×10^-9^M compared to 7.1×10^-11^M, respectively (Figure 1B). These inhibitory effects were mediated via CGRP-R activation, as pre-incubation with the CGRP-R antagonist BIBN4096 completely abrogated CGRP- and SAX-mediated inhibition (Figure 1C). We previously showed that one of the functional effects of CGRP in LCs is to increase expression of the HIV-1-binding LCs-specific C-type lectin langerin (7–9). Accordingly, CGRP and SAX increased langerin surface expression in MDLCs in a dose-dependent manner, with SAX having lower potency (Figure 1D), as for trans-infection inhibition.

**Figure 1.**
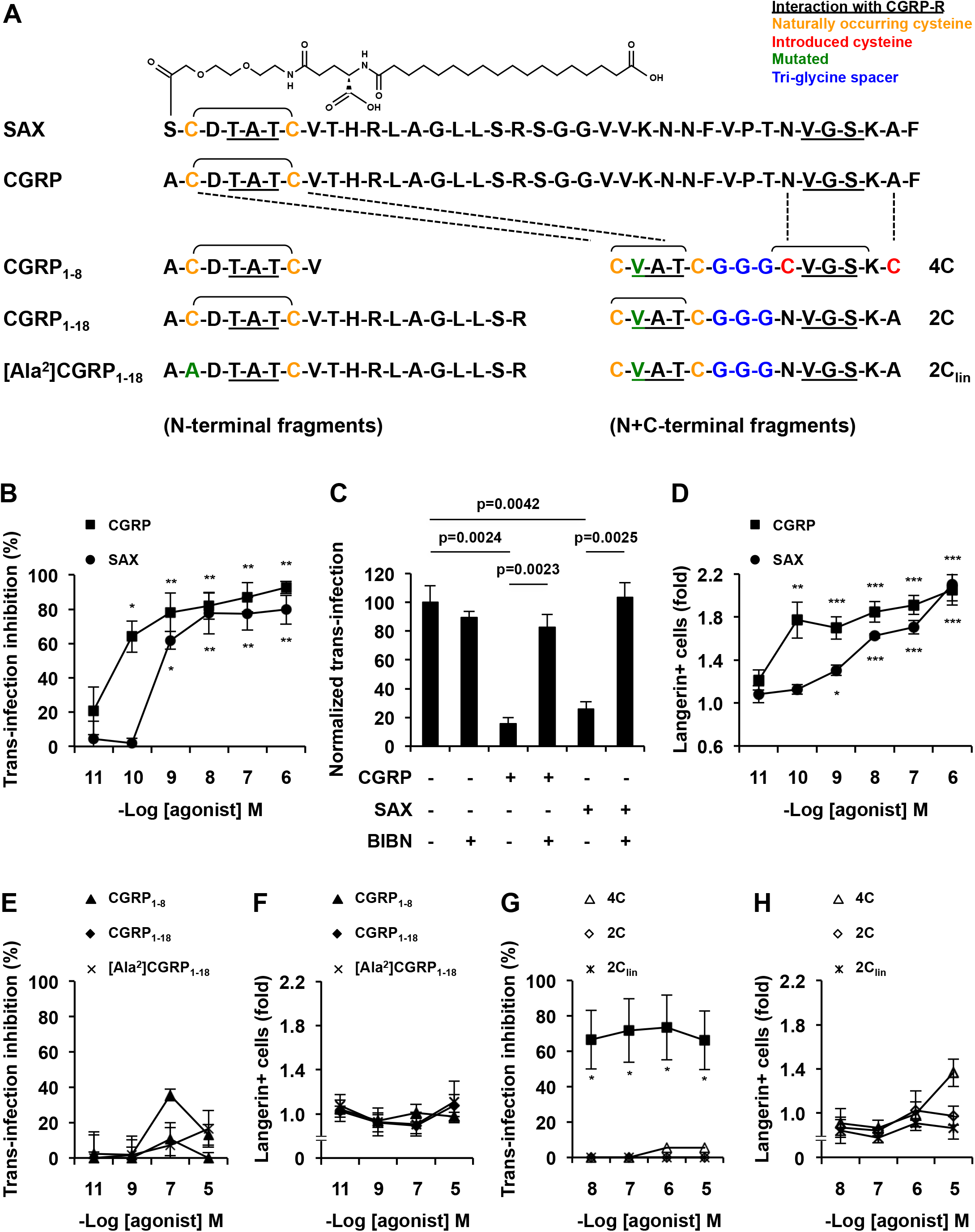
CGRP and SAX, but not CGRP peptide fragments, inhibit LCs-mediated HIV-1 trans-infection and increase langerin expression. **(A)** Sequences of CGRP, SAX, and the different N/N+C-terminal fragments used. **(B-H)** MDLCs were left untreated or treated for 24h with the indicated concentrations of CGRP, SAX or CGRP peptide fragments. When indicated, the CGRP-R antagonist BIBN4096 (BIBN, 1μM) was added 15min before addition of agonists. The cells were then pulsed with HIV-1 JRCSF for 2h, washed, and incubated for 7 days with autologous CD4+ T-cells (B, C, E) or 3 days with HIV-1 GFP-reporter CCR5^high^CD4^high^ T-cells (G). HIV-1 trans-infection and replication in T-cells was evaluated by p24 ELISA in the co-culture supernatants or by GFP expression and flow cytometry. In other experiments, the cells were stained for langerin surface expression and examined by flow cytometry (D, F, H). Shown are means±SEM (n≥3) of HIV-1 trans-infection inhibition (B, E, G) or normalized (C), and folds increase in langerin expression (D, F, H); p values are derived from two-sided Student’s t-test.

In contrast, the CGRP N-terminal fragments CGRP_1-8_ (11), CGRP_1-18_ (12) and the negative control mutated [Ala^2^]CGRP_1-18_ lacked agonistic activity and did not significantly inhibit LCs-mediated trans-infection (Figure 1E). Similarly, none of these CGRP N-terminal fragments increased langerin surface expression (Figure 1F). We also designed novel bivalent CGRP fragments (Figure 1A) by linking with a triglycine spacer the previously described constrained CGRP N- and C-terminal regions (12), containing disulphide bonds either at both N/C-terminal regions or only in the N-terminal. None of these bivalent CGRP peptide fragments significantly inhibited LCs-mediated HIV-1 trans-infection (Figure 1G) or increased langerin surface expression (Figure 1H).

These results show that in order to inhibit LCs-mediated HIV-1 trans-infection, CGRP-R activation requires full-length CGRP or SAX. We speculate that the lack of activity by CGRP peptide fragments might be related to their ‘biased signaling’ (13) across CGRP-R, which alike other G-protein coupled receptors activates multiple downstream signaling pathways. Hence, compared to the native CGRP ligand, CGRP fragments might have allosteric bias for preferential activation of particular pathways, which are not the ones mediating inhibition of HIV-1 trans-infection, i.e. while CGRP activates its canonical cAMP/PKA signaling in LCs (14), we showed that the anti-HIV-1 effects of CGRP in LCs are mediated via NFκB/STAT4 signaling (8). Alternatively, further optimization of the CGRP peptide fragments might be required. For instance, the N+C-terminal fragments could be re-designed to better fit into CGRP-R binding pockets with higher affinity, potentially including longer/different spacer regions.

### CGRP limits mucosal HIV-1 transmission *in-vivo*

We investigated the potential inhibitory effects of topically applied CGRP in three different *ex-vivo* and/or *in-vivo* mucosal models. First, we prepared polarized human penile tissue explants, as we previously described (15), from the stratified and nonkeratinized *fossa navicularis* region that structurally resembles the vaginal epithelium. Topical application of biotinylated CGRP onto the apical side for 3h, followed by histochemistry, showed that CGRP readily penetrated the mucosal epithelial, but not the stromal compartment (Figure 2A). Second, as CGRP mediates vasodilator-dependent neurogenic inflammation that can result in immune cells recruitment, we topical applied CGRP onto the vagina of normal BALB/c mice for 6h. We found that CGRP was not toxic, did not induce overt inflammation, and did not modify the distribution and density of T-cells, B-cells and macrophages (Figure 2B). Third, we used bone-marrow/liver/thymus (BLT) humanized mice, in which we confirmed the reconstitution and presence of human langerin-expressing LCs within the vaginal epithelial compartment (Figure 2C). Single topical application of CGRP onto the vagina of BLT mice for 4h, followed by a vaginal challenge with high dose HIV-1 (Figure 2D), dose-dependently restricted the increase in plasma viral load (Figure 2E) and maintained higher CD4+ T-cell counts (Figure 2F).

**Figure 2.**
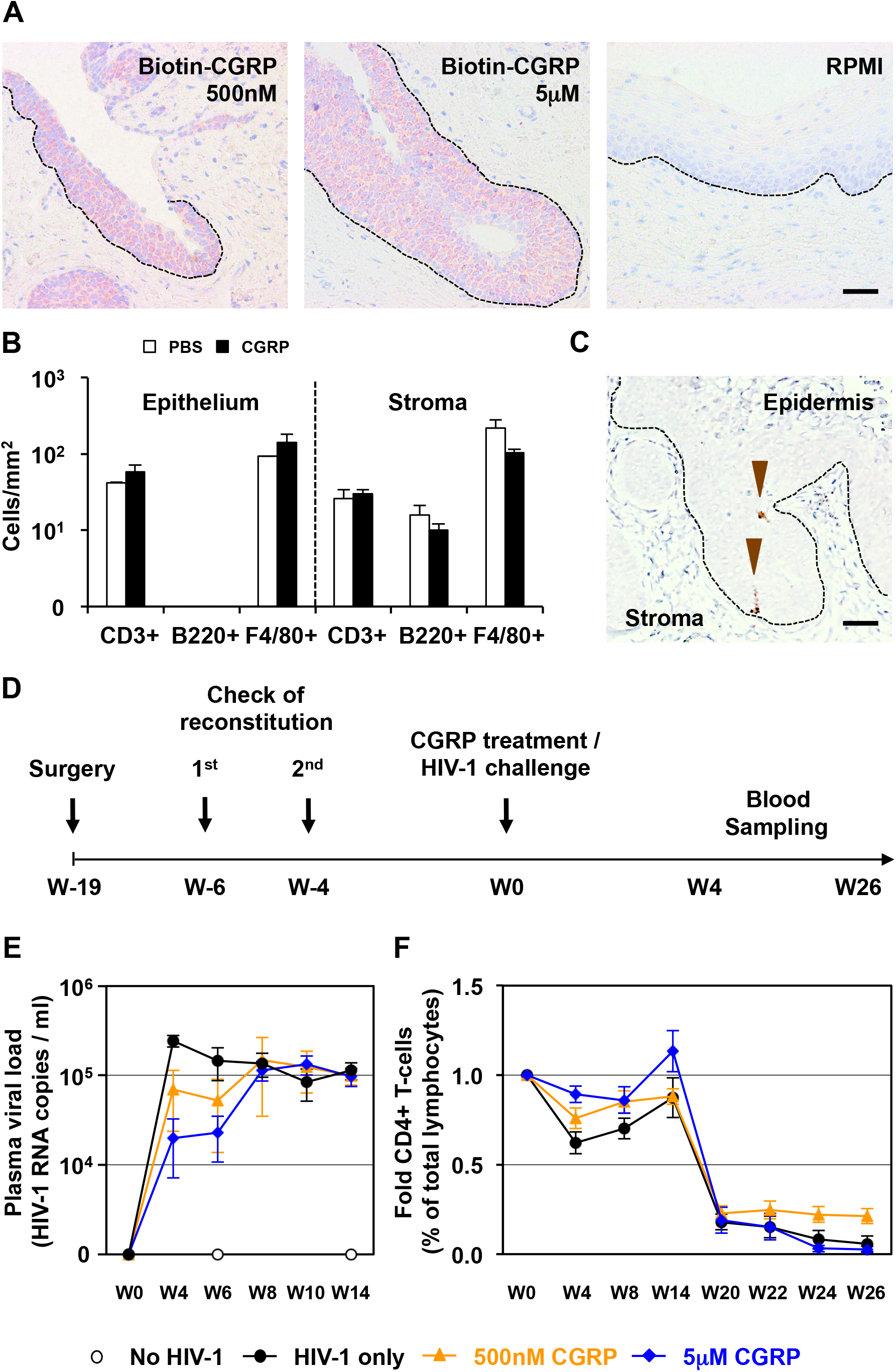
CGRP inhibits HIV-1 transmission *in-vivo*. **(A)** Entry of biotinylated CGRP into *fossa navicularis* explants, revealed with streptavidin-HRP, AEC peroxidase substrate (red), and hematoxyline counterstaining (blue). Images are representative of n=3 tissues; broken lines denote the basement membranes and scale bar = 20μm. **(B)** CGRP (1μM) or PBS were topically applied intravaginally in normal female BALB/c mice. Vaginal tissue sections were examined by immunohistochemistry for the presence of CD3+ T-cells, B220+ B-cells and F4/80+ macrophages. Numbers represents mean±SEM cells/mm^2^ in either the vaginal epithelium or stroma. **(C)** Representative image of the vaginal tissue of humanized BLT mice, showing expression of human langerin (arrowheads); scale bar = 20μm. **(D)** Experimental schedule for preparing humanized female BLT mice, intravaginally applying CGRP or PBS followed by challenge with high-dose HIV-1 JRCSF, and subsequent blood sampling for measurement of plasma viral loads and T-cell counts. **(E, F)** Mean plasma viral loads (HIV-1 RNA copies / ml) or CD4+ T-cell counts (percentages of total lymphocytes) in BLT mice (n=5 animals per group).

As T-cells recruitment was absent in CGRP-treated mice, we speculate that CD4+ T-cell maintenance is mediated by CGRP acting on vaginal LCs and inhibiting their capacity to disseminate HIV-1 to CD4+ T-cells. Indeed, CGRP entry into mucosal tissues was restricted to the epithelial compartment, in which LCs reside. Moreover, we previously showed that CGRP treatment of CD4+ T-cells, rather than LCs, has no effect on HIV-1 trans-infection (7). We further speculate that in order to achieve better and long-lasting viremia and CD4+ T-cell control, CGRP treatment should probably be longer, with repeated and continuous applications similar to pre-exposure prophylactic (PreP) therapy, and that a high potency mucosal metabolically stable analogue would be beneficial.

Collectively, our findings provide proof-of-concept that CGRP-based interventions, namely high potency mucosal metabolically stable analogues and/or optimized agonists, could potentially be used for the prophylactic prevention of mucosal HIV-1 transmission. As such, HIV-1 infection should be included within the different pathologies and inflammatory conditions, in which CGRP is beneficial and could be harnessed to exert protective effects.

## Materials and methods

### *In-vitro* experiments

We used the following materials: CGRP and BIBN4096 (Sigma), SAX (prepared as previously described (2)) and custom synthesized CGRP peptide fragments (United Biosystems). Langerin surface expression by flow cytometry and HIV-1 transinfection, using MDLCs, autologous CD4+ T-cells or green fluorescent protein (GFP)-reporter T-cells, were performed as we described (7–9). IC_50_ values were calculated with Prism software (GraphPad) using the log(inhibitor) vs. normalized response - variable slope model.

### *Ex-vivo* experiments

We used human penile tissues obtained under informed consent and ethical approval, as part of our previous study (15). Polarized *fossa navicularis* explants were prepared as we described (15), and exposed for 3h to 500nM or 5μM biotinylated CGRP (AnaSpec) in 100μl RPMI medium added to the apical side. After incubation, tissue entry of biotinylated CGRP was examined by histochemistry of 4μm paraffin sections as we described (15), using streptavidin coupled to horseradish peroxidase (HRP; Vector), followed by the red 3-amino-9-ethylcarbazole (AEC) HRP substrate (Dako) and counterstaining with haematoxylin. Images were acquired with an Olympus BX63F microscope using MetaMorph (Molecular Devices) and analyzed with ImageJ software (NIH).

### *In-vivo* experiments

We used female BALB/c mice (10 weeks old, 25-30gr, three mice per group, synchronized in estrous cycle, n=3 experiments), under ethical approval from the institutional review board of the San Raffaele Scientific Institute (IACUC no. 599). CGRP, at 10nM, 100nM or 1μM, was diluted in 30μl sterile PBS, alone or in combination with 1% hydrocortisone, and applied intravaginally for 6h. Spleen, lymph nodes, gut, liver, kidneys and female reproductive system were then collected, and haematoxylin and eosin stained 3μm paraffin sections were examined for histopathological analysis. Selected slides were stained after antigen retrieval with monoclonal antibodies (Bio-Rd), including rat-anti-human CD3, rat-anti-mouse B220/CD45R (clone RA3-6B2) and rat-anti-mouse F4/80 (clone CI:A31), followed by rat on rodent HRP-polymer, 3,3’-diaminobenzidine (DAB) as chromogen (Biocare Medical) and counterstaining with haematoxylin. Images were acquired with the AxioCam HRc using the AxioVision System SE64 (Zeiss).

Humanized female BLT mice were prepared at the Ragon Institute Human Immune System Mouse Program, according to their established protocols (see https://www.ragoninstitute.org/research/services/humanized-mouse/). Expression of human langerin was determined by immunohistochemistry of 4μm vaginal tissue paraffin sections as we described, using goat-anti-human langerin Ab (R&D), the LSAB2 System-HRP with the brown di-amino benzidine (DAB) substrate (Dako) and counterstaining with haematoxylin. CGRP, at 500nM or 5μM, in 30μl sterile PBS or PBS alone were topically applied (five mice per group) onto the vaginal epithelium for 4h, followed by topical vaginal challenge with 2×10^4^ TCID_50_ HIV-1 JRCSF (NIH, prepared as before (6).

## Acknowledgments

The study was supported by grants from ANRS (France Recherche Nord & sud Sida-hiv hépatites) and La SATT IDFinnov.

## Notes

### Competing Interest Statement

The authors have declared no competing interest.

